# Target-directed microRNA degradation regulates developmental microRNA expression and embryonic growth in mammals

**DOI:** 10.1101/2023.06.26.546601

**Authors:** Benjamin T Jones, Jaeil Han, He Zhang, Robert E. Hammer, Bret M. Evers, Dinesh Rakheja, Asha Acharya, Joshua T. Mendell

## Abstract

MicroRNAs (miRNAs) are post-transcriptional regulators of gene expression that play critical roles in development and disease. Target-directed miRNA degradation (TDMD), a pathway in which miRNAs that bind to specialized targets with extensive complementarity are rapidly decayed, has emerged as a potent mechanism of controlling miRNA levels. Nevertheless, the biological role and scope of miRNA regulation by TDMD in mammals remains poorly understood. To address these questions, we generated mice with constitutive or conditional deletion of *Zswim8*, which encodes an essential TDMD factor. Loss of *Zswim8* resulted in developmental defects in heart and lung, growth restriction, and perinatal lethality. Small RNA sequencing of embryonic tissues revealed widespread miRNA regulation by TDMD and greatly expanded the known catalog of miRNAs regulated by this pathway. These experiments also uncovered novel features of TDMD-regulated miRNAs, including their enrichment in co-transcribed clusters and examples in which TDMD underlies ‘arm switching’, a phenomenon wherein the dominant strand of a miRNA precursor changes in different tissues or conditions. Importantly, deletion of two miRNAs, miR-322 and miR-503, rescued growth of *Zswim8* null embryos, directly implicating the TDMD pathway as a regulator of mammalian body size. These data illuminate the broad landscape and developmental role of TDMD in mammals.

## Introduction

MicroRNAs (miRNAs) are ∼22 nucleotide RNAs that negatively regulate messenger RNA (mRNA) stability and translation (Bartel 2018). miRNAs act as obligate co-factors for Argonaute (AGO) proteins, which they guide to target mRNAs primarily through base-pairing interactions between the miRNA 5’ end, termed the seed sequence, and complementary sites that are most often located in the 3¢ untranslated regions (UTRs) of targets. Binding of AGO proteins results in recruitment of deadenylation and decapping complexes, leading to target repression (Jonas and Izaurralde 2015). Studies over the last two decades have established that miRNA-mediated regulation is critical for development and physiology in diverse metazoan species (Vidigal and Ventura 2015; Bartel 2018). Accordingly, elaborate mechanisms that impact miRNA abundance and activity have evolved to precisely control how miRNAs are deployed to regulate gene expression (Gebert and MacRae 2019).

Once loaded into an AGO protein, the 5¢ and 3¢ termini of the miRNA are deeply buried in the AGO middle (mid) and PIWI/AGO/Zwille (PAZ) domains, respectively, thereby protecting the miRNA from exonucleolytic decay (Elkayam et al. 2012; Schirle and MacRae 2012). As a result, miRNAs are typically extremely stable, with half-lives extending to days or weeks in vivo (van Rooij et al. 2007; Gatfield et al. 2009; Kingston and Bartel 2019). Nevertheless, the existence of miRNAs with accelerated decay rates, first observed in mammalian cell lines (Hwang et al. 2007; Bail et al. 2010; Krol et al. 2010; Gantier et al. 2011; Rissland et al. 2011; Guo et al. 2015; Marzi et al. 2016; Kingston and Bartel 2019; Reichholf et al. 2019), foreshadowed the discovery of mechanisms that carry out sequence-specific miRNA degradation. Of the reported mechanisms of accelerated miRNA turnover, the most extensively studied is target-directed miRNA degradation (TDMD). TDMD is triggered when a miRNA engages specialized target sites with extended complementarity to both the seed sequence and the miRNA 3¢ end. This miRNA decay pathway was initially discovered as a mechanism used by viruses to remodel host miRNA expression (Cazalla et al. 2010; Libri et al. 2012; Marcinowski et al. 2012; Lee et al. 2013) and, simultaneously, was shown to be induced by expression of highly complementary synthetic miRNA targets in *Drosophila* and human cells (Ameres et al. 2010). The subsequent identification of endogenous mammalian transcripts that function as triggers for TDMD, specifically *Cyrano*, *Nrep*, and *Serpine1*, which promote decay of miR-7, miR-29b, and miR-30b/c, respectively, established that this pathway is employed as a natural mechanism to regulate the abundance of specific miRNAs in vivo (Bitetti et al. 2018; Ghini et al. 2018; Kleaveland et al. 2018).

Multiple studies recently reported the discovery of a cullin-RING ubiquitin ligase (CRL) complex containing the substrate adapter protein ZSWIM8 as a key mediator of TDMD (Han et al. 2020; Shi et al. 2020). The extensive complementarity between miRNAs and TDMD-inducing targets results in a conformational change in AGO (Sheu-Gruttadauria et al. 2019) that is believed to be specifically recognized by the ZSWIM8 complex. This results in AGO ubiquitylation, degradation by the proteasome, and consequent release of the miRNA for decay by cytoplasmic nucleases. When engaged with TDMD-inducing targets, the extended base-pairing of the miRNA 3¢ end also results in its release from the AGO PAZ domain (Sheu-Gruttadauria et al. 2019). This renders it susceptible to the activity of polymerases that add non-templated nucleotides or exoribonucleases that remove nucleotides, a process termed tailing and trimming (Ameres et al. 2010). Although tailing and trimming is not essential for the activity of the ZSWIM8 complex or degradation of the miRNA (Han et al. 2020; Shi et al. 2020), its strong correlation with the TDMD pathway has proven to be a useful feature for identifying TDMD-inducing target transcripts (Li et al. 2021).

These advances in our understanding of the mechanism of TDMD have enabled broader investigation of the landscape of miRNAs that are regulated by this pathway in model organisms and cell lines. Indeed, loss-of-function of ZSWIM8 or its orthologs (EBAX-1 in *C. elegans*, Dora in *Drosophila*), coupled with small RNA sequencing, has revealed dozens of miRNAs that are strongly regulated by TDMD in mammalian cell lines (Han et al. 2020; Shi et al. 2020), *Drosophila* cell lines and embryos (Shi et al. 2020; Kingston et al. 2022), and adult *C. elegans* (Shi et al. 2020). For a subset of the newly-identified mammalian and *Drosophila* TDMD-regulated miRNAs, decay-inducing target sites in both coding and noncoding transcripts that exhibit complementarity to both the miRNA seed sequence and 3¢ end have been identified (Li et al. 2021; Kingston et al. 2022; Sheng et al. 2023). Interestingly, in *C. elegans*, miRNAs belonging to the *miR-35* family are degraded at the embryo to L1 transition in an EBAX-1-dependent manner, suggesting that they are additional substrates of the TDMD pathway (Donnelly et al. 2022). Nevertheless, degradation of these miRNAs relies only on their seed sequences, without a requirement for 3¢ complementarity, suggesting the existence of alternative modes of recruiting ZSWIM8 orthologs for decay of specific miRNAs in worms and possibly other species.

The discovery of the ZSWIM8 complex has also allowed exploration of the biological roles of TDMD in animals. Dysregulation of miRNAs in Dora-mutant *Drosophila* results in embryonic lethality and abnormal cuticle development, establishing an essential role for TDMD in development in flies (Kingston et al. 2022). Disentangling the developmental functions of ZSWIM8 homologs in *C. elegans* and mammals, however, has proven more challenging. Worms lacking EBAX-1 are viable and morphologically normal, but exhibit an axon guidance defect that has been attributed to EBAX-1-mediated degradation of misfolded SAX-3, a receptor required for axonal pathfinding (Wang et al. 2013). Conditional deletion of *Zswim8* in mouse brain leads to incompletely penetrant perinatal lethality, reduced body size, and widespread neurodevelopmental abnormalities (Wang et al. 2023). These defects were reported to result, at least in part, from loss of ZSWIM8-mediated degradation of misfolded DAB1, a factor with pleiotropic functions in neuronal development. Thus, the role of TDMD in mammalian development and physiology, and the landscape of mammalian miRNAs regulated by this pathway in vivo, remains to be determined.

To address these outstanding questions, we generated mice harboring germline-transmitted and conditional *Zswim8* loss-of-function alleles. We observed several highly-penetrant phenotypes in *Zswim8^‒/‒^* mice, including growth restriction in the developing embryo, defects in heart and lung development, and perinatal lethality. Small RNA sequencing of late embryonic tissues revealed widespread regulation of miRNAs by TDMD in vivo and greatly expanded the known set of miRNAs regulated by this pathway in mammals. Importantly, we demonstrated that stabilization of two co-transcribed miRNAs, miR-322 and miR-503, plays a causative role in growth restriction of *Zswim8^‒/‒^* embryos, thus establishing the TDMD pathway as a potent regulator of body size during mammalian development.

## Results

### Loss of ZSWIM8 causes perinatal lethality and developmental defects in mice

To investigate the biological role of TDMD in mammals, we used CRISPR/Cas9 genome editing of mouse zygotes to generate multiple constitutive and conditional *Zswim8* loss-of-function alleles. To achieve global loss-of-function, dual single guide RNAs (sgRNAs) were used to delete *Zswim8* exons 2 through 7, which encode the BC-box and Cullin-2-box necessary for formation of the ZSWIM8 CRL complex (Wang et al. 2013), as well as the conserved SWIM domain (**Fig. 1A**). Loss of these exons is also predicted to result in a frameshift mutation that introduces a premature termination codon. Mice homozygous for this allele are hereafter designated *Zswim8*^‒/‒^. Additionally, we generated a conditional knockout allele by introducing loxP sites flanking exon 2 (**Fig. 1A**). Loss of this exon removes the BC- and Cullin-2 boxes and disrupts the *Zswim8* reading frame. Animals homozygous for this allele are referred to as *Zswim8*^F/F^.

**Figure 1.**
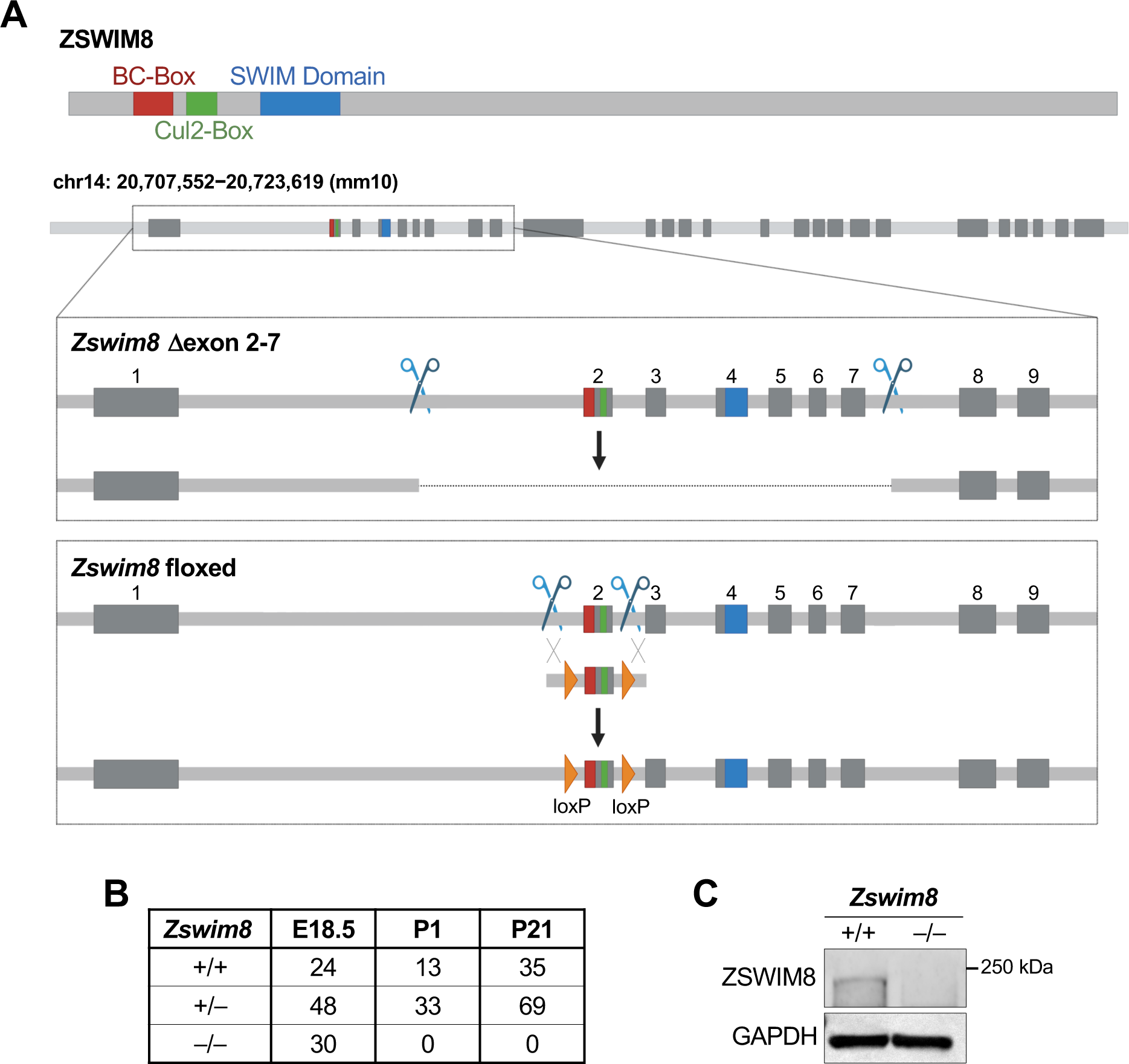
Loss of ZSWIM8 results in perinatal lethality in mice. (*A*) Genome editing strategy used to generate *Zswim8*^‒/‒^ (ι1exon 2-7) and *Zswim8*^F/F^ (floxed) mice. Upper, schematic of ZSWIM8 protein with approximate locations of BC-box (red), Cul2-box (green), and SWIM domain (blue) indicated. Lower, depiction of mouse *Zswim8* genomic locus with exons encoding each domain in appropriate color as defined in upper panel, scissors showing CRISPR-Cas9 targeting sites, and DNA segment with orange triangles indicating donor sequence containing loxP sites. (*B*) Frequency of genotypes of offspring produced from *Zswim8*^+/‒^ intercrosses at the indicated time-points. (*C*) Western blot of ZSWIM8 protein in E18.5 brain from embryos of the indicated genotypes.

Mice heterozygous for the exon 2-7 deletion were intercrossed to determine the consequences of global ZSWIM8 loss of function. Whereas wild-type and heterozygous animals were present at the expected frequencies at weaning (post-natal day 21; P21), no *Zswim8*^‒/‒^ mice were observed at this time-point (**Fig. 1B**). To determine at which stage *Zswim8*^‒/‒^ mice were lost, timed matings were performed and litters were delivered by cesarean section at embryonic day 18.5 (E18.5). At this stage, all genotypes were observed at the expected Mendelian frequencies (**Fig. 1B**). We noted, however, that *Zswim8*^‒/‒^ pups exhibited agonal breathing and died shortly after delivery. Furthermore, no *Zswim8*^‒/‒^ mice survived to P1 (**Fig. 1B**). Thus, loss of *Zswim8* results in perinatal lethality in mice.

To determine the cause of perinatal death, we further examined *Zswim8*^‒/‒^ embryos at E18.5. Western blotting confirmed the expected loss of ZSWIM8 protein in knockout tissues (**Fig. 1C**). *Zswim8*^‒/‒^ embryos were significantly smaller than their wild-type and heterozygote littermates, with a >20% reduction in overall body weight (**Fig. 2A-B**). A complete necropsy of *Zswim8*^‒/‒^ embryos further revealed overt and highly-penetrant defects in heart morphology at this time-point. Hearts from knockout animals were grossly smaller and globose in shape, lacking a distinct apex (**Fig. 2C**). Histologic analysis revealed dilated ventricular cavities with thinning and non-compaction of ventricular walls (**Fig. 2D**). We additionally observed ventricular septal defects (VSDs) in approximately one-third of *Zswim8^‒/‒^* hearts (**Supplemental Fig. S1A**).

**Figure 2.**
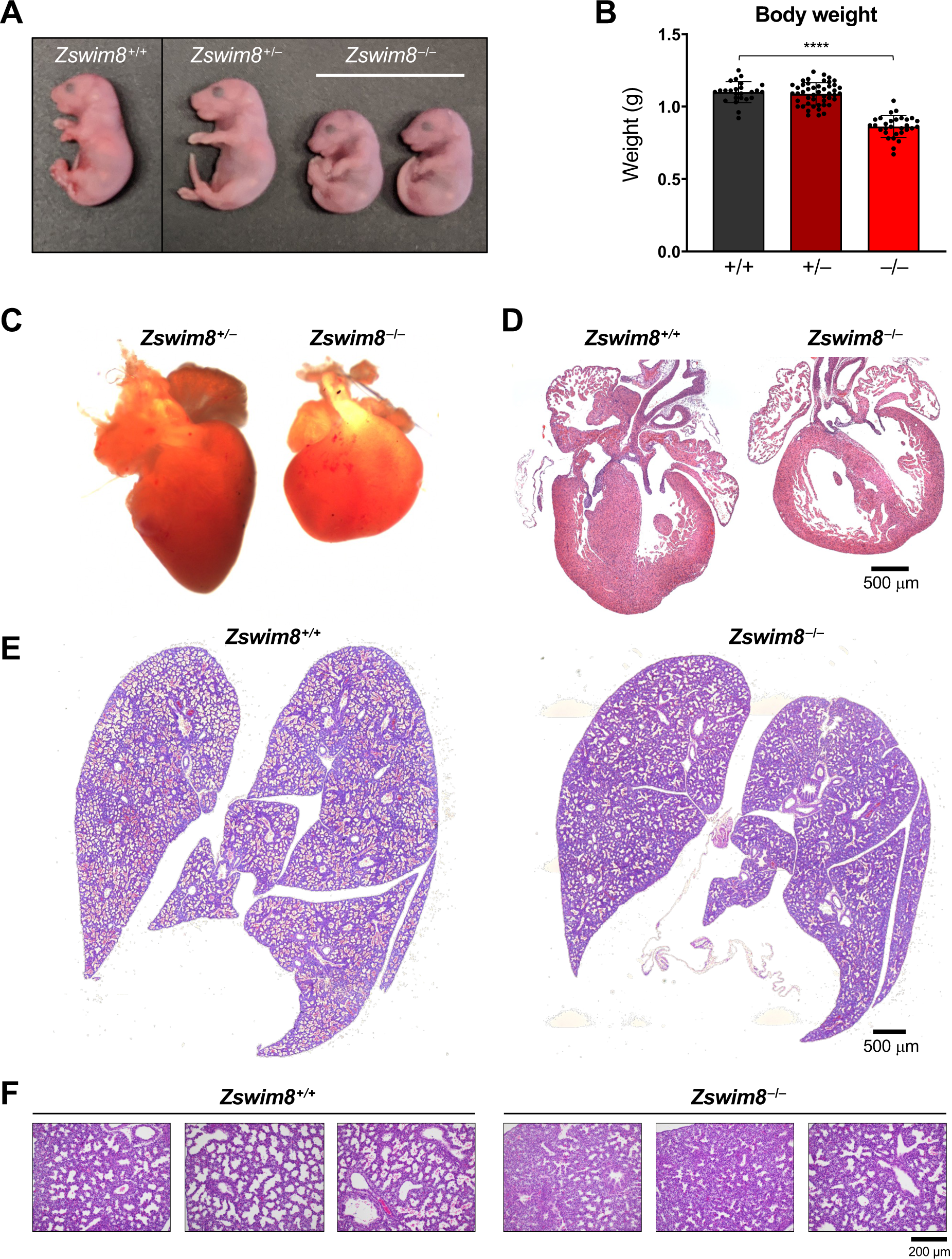
ZSWIM8 deficiency leads to growth restriction and defective heart and lung development. (*A*) Images of E18.5 mice of the indicated genotypes. (*B*) Body weights of E18.5 mice of the indicated genotypes. n=25 (*Zswim8*^+/+^), 47 (*Zswim8*^+/‒^), and 30 (*Zswim8*^‒/‒^). Data are represented as mean ± SD with individual data points shown. ****p<0.0001 (unpaired *t*-test). (*C*) Images of E18.5 hearts of the indicated genotypes. (*D*-*F*) Representative hematoxylin and eosin (H&E)-stained sections of E18.5 hearts (*D*) and lungs (*E*-*F*) of the indicated genotypes.

To determine if abnormal cardiac development was the cause of perinatal lethality, we crossed mice carrying the floxed *Zswim8* allele to mice harboring a *Nkx2.5*-Cre transgene that is expressed as early as E7.5 in the cardiac crescent and in the majority of heart tube progenitor cells (McFadden et al. 2005). Although we observed a slight under-transmission of the *Nkx2.5*-Cre transgene overall, *Nkx2.5*-Cre; Z*swim8*^+/+^ and *Nkx2.5*-Cre; Z*swim8*^F/F^ mice were equally represented at P21 (**Supplemental Fig. S1B**). Moreover, the body weight of *Nkx2.5*-Cre; Z*swim8*^F/F^ mice at E18.5 was equivalent to control animals (**Supplemental Fig. S1C**). Notably, *Nkx2.5*-Cre; Z*swim8*^F/F^ mice exhibited developmental heart defects that resembled those observed in *Zswim8*^‒/‒^ embryos, including less defined apices and dilation of the ventricles, although these phenotypes were less severe than those observed in global knockout animals (**Supplemental Fig. S1D)**. Quantitative PCR documented successful recombination of the floxed allele in *Nkx2.5*-Cre; Z*swim8*^F/F^ hearts (**Supplemental Fig. S1E**). Although a significant fraction of detectable alleles remained unrecombined, this was likely due in part to a contribution from non-cardiac lineages. Overall, these data demonstrated that intrinsic deficiency of ZSWIM8 in the embryonic heart impairs cardiac development, but these abnormalities may not account for perinatal lethality of *Zswim8*^‒/‒^ mice. It remains possible, however, that earlier and/or complete loss of the protein in developing hearts of germline *Zswim8* knockout animals produces more severe cardiac defects that are incompatible with post-natal viability.

Seeking other potential causes neonatal death of *Zswim8*^‒/‒^ mice, we next examined the lungs of these animals. Defects in lung development are a common cause of perinatal lethality (Turgeon and Meloche 2009) and were of particular interest due to the agonal breathing displayed by newborn *Zswim8*^‒/‒^ mice. At E16.5, mouse lungs enter the canalicular stage of development, in which the respiratory tree expands and is vascularized (Warburton et al. 2010). This is followed by the saccular stage, extending from E17.5 to P5, during which thinning of the septa occurs, alveolar airspaces expand, and epithelial cells differentiate into the alveolar type I and type II pneumocytes to produce surfactant. Lungs were harvested from E18.5 *Zswim8*^+/+^ and *Zswim8*^‒/‒^ mice before animals initiated breathing. As expected, wild-type lungs displayed features characteristic of the mid-to-late saccular stage of development, with emerging airspaces and thinning alveolar septa (**Fig. 2E-F**). Lungs from *Zswim8*^‒/‒^ embryos, in contrast, appeared less mature, with less expanded saccules, thicker septa, and minimally expanded terminal airspaces. These findings were indicative of the late canalicular stage of development. Thus, ZSWIM8-deficiency resulted in delayed lung development, providing a likely explanation for respiratory failure in newborn *Zswim8*^‒/‒^ mice. Altogether, these results established an essential role for ZSWIM8 in mammalian development, with critical roles in organogenesis of the heart and lung and in regulation of body size.

### The landscape of TDMD-regulated miRNAs in mammalian tissues

We hypothesized that the stabilization of miRNAs that are normally degraded through the TDMD pathway contributes to the developmental defects observed in *Zswim8*^‒/‒^ mice. To identify such miRNAs, and to comprehensively characterize the landscape of mammalian miRNAs that are regulated by TDMD in vivo, we performed small RNA sequencing on a broad panel of tissues from E18.5 *Zswim8*^+/+^ and *Zswim8*^‒/‒^ mice (brain, heart, lung, liver, small intestine, kidney, skin, and stomach). These studies revealed numerous miRNAs that were strongly upregulated in *Zswim8*^‒/‒^ tissues (**Fig. 3A** and **Supplemental Table S1**). To distinguish between miRNAs that were upregulated due to increased transcription or processing, possibly as a secondary consequence of the developmental abnormalities in ZSWIM8-deficient tissues, from those whose stability was increased due to abrogation of TDMD, we examined both strands of each miRNA duplex. TDMD occurs after the loading of the mature miRNA into AGO (Han et al. 2020; Shi et al. 2020), which separates it from the opposite strand of the processed miRNA duplex, also known as the passenger strand or miRNA*. As a consequence, direct TDMD substrates are expected to be upregulated upon loss of ZSWIM8 without an increase in abundance of the opposite strand. On the basis of this criterion, we detected 57 miRNAs that behaved as direct substrates of the TDMD pathway in at least one tissue, thereby nearly doubling the known number of miRNAs regulated by TDMD in mammals (**Fig. 3B**). While many miRNAs were broadly controlled by TDMD across all sequenced tissues, some were regulated in a tissue-specific manner, suggesting the restricted expression of the target transcripts that induce their decay. Northern blotting of a selected panel of miRNAs confirmed their robust upregulation in ZSWIM8-deficient tissues (**Fig. 3C** and **Supplemental Fig. S2**).

**Figure 3.**
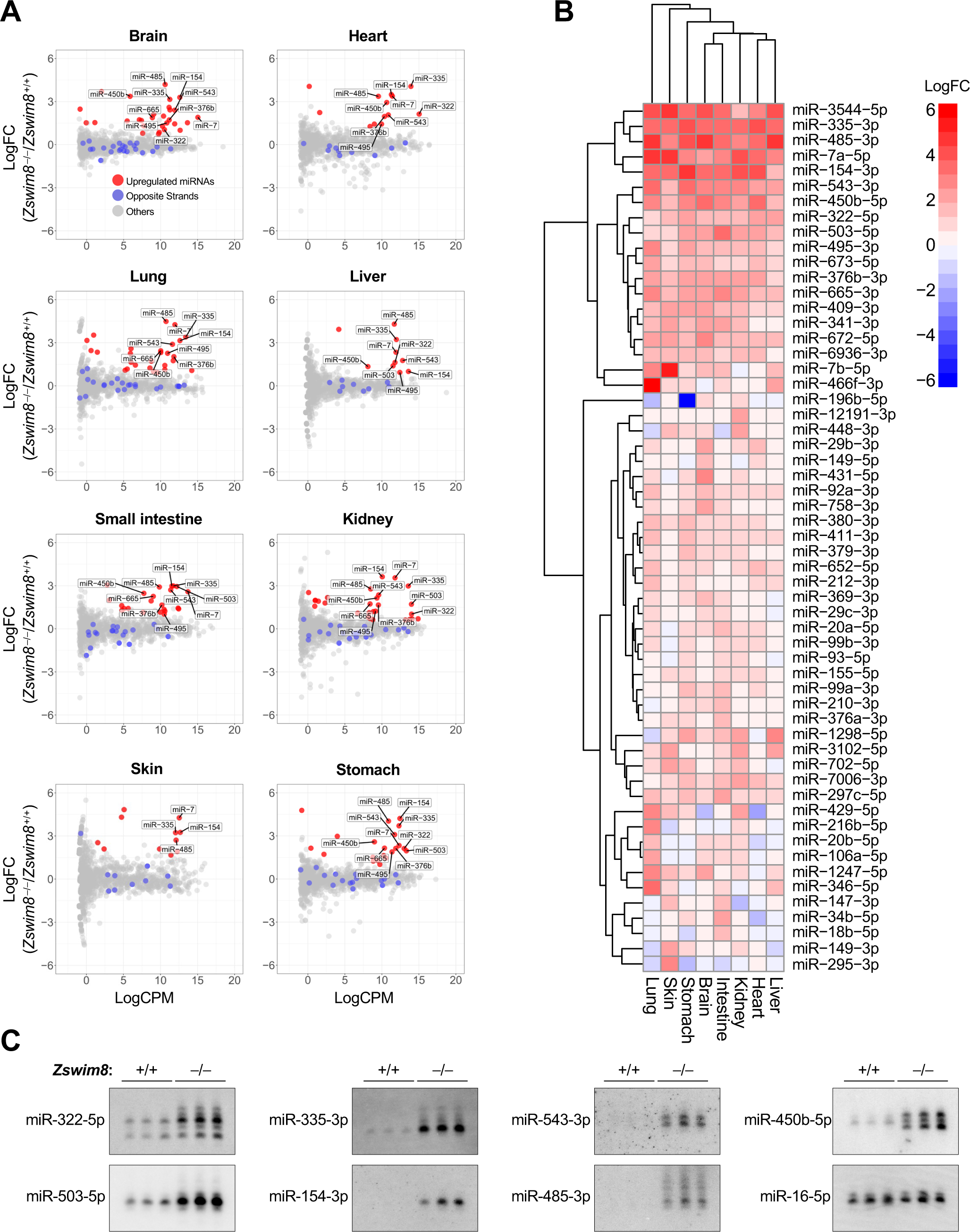
The landscape of TDMD-regulated miRNAs in mouse embryonic tissues. (*A*) Small RNA sequencing of E18.5 tissues. Red dots show miRNAs that were significantly upregulated in *Zswim8*^‒/‒^ relative to *Zswim8*^+/+^ tissues (p<0.05; FDR<0.05) without a corresponding increase in the opposite strand derived from the same precursor (labeled as blue dots). n=3 biological replicates per genotype. LogFC, log_2_ fold change; LogCPM, log_2_ counts per million in *Zswim8*^+/+^ tissue. (*B*) Heat map showing log_2_ fold change of all TDMD-regulated miRNAs (*Zswim8*^‒/‒^/*Zswim8*^+/+^), defined as those exhibiting significant upregulation in at least one *Zswim8*^‒/‒^ tissue (p<0.05; FDR<0.05) without a corresponding increase in the opposite strand derived from the same precursor. (*C*) Northern blot analysis of miRNA expression in E18.5 hearts from mice of the indicated genotypes. n=3 biological replicates per genotype. miR-16-5p served as a loading control.

During the loading of miRNAs into AGO, the strands of the processed miRNA duplex are discriminated such that loading of the mature miRNA, or guide strand, is favored, while the dissociated passenger strand is discarded and degraded (Kim et al. 2009). Consequently, the more abundant strand is generally defined as the guide and the less abundant strand is designated the passenger, although examples in which both strands are loaded and regulate targets have been described (Medley et al. 2021). We observed that 12 of the 57 ZSWIM8-regulated miRNAs represented the normally less abundant strand of the miRNA duplex (miR-99a-3p, miR-99b-3p, miR-149-3p, miR-154-3p, miR-335-3p, miR-379-3p, miR-411-3p, miR-429-5p, miR-652-5p, miR-702-5p, miR-3102-5p, miR-3544-5p), suggesting that these miRNAs are loaded into AGO and actively engage targets despite being annotated as passenger strands. For four of these miRNAs (miR-154, miR-335, miR-411, miR-3544), the normally less abundant strand became the more abundant strand in one or more ZSWIM8-deficient tissues (**Fig. 4A**). The phenomenon in which the dominant strand of a miRNA duplex changes in different cell-types or conditions has been referred to as ‘arm-switching’, and is generally believed to be a mechanism for regulating miRNA activity that operates at the AGO loading step (Grimson et al. 2008; Marco et al. 2010; Cloonan et al. 2011; Griffiths-Jones et al. 2011; Li et al. 2011; Li et al. 2012). Our data suggested that some examples of arm switching may be a consequence of differential regulation of specific miRNA strands by the TDMD pathway. For example, miR-154, a miRNA that has been reported to undergo arm switching (Cheng et al. 2013), exhibits 3 prime (3p) dominance in brain, intestine, liver, and lung, while exhibiting 5p dominance in heart, kidney, skin, and stomach (**Fig. 4A**). In *Zswim8*^‒/‒^ mice, however, all tissues exhibit a strong miR-154-3p bias, suggesting that differential regulation of the 3p strand by TDMD across tissues most likely accounts for the apparent arm switching of this miRNA.

**Figure 4.**
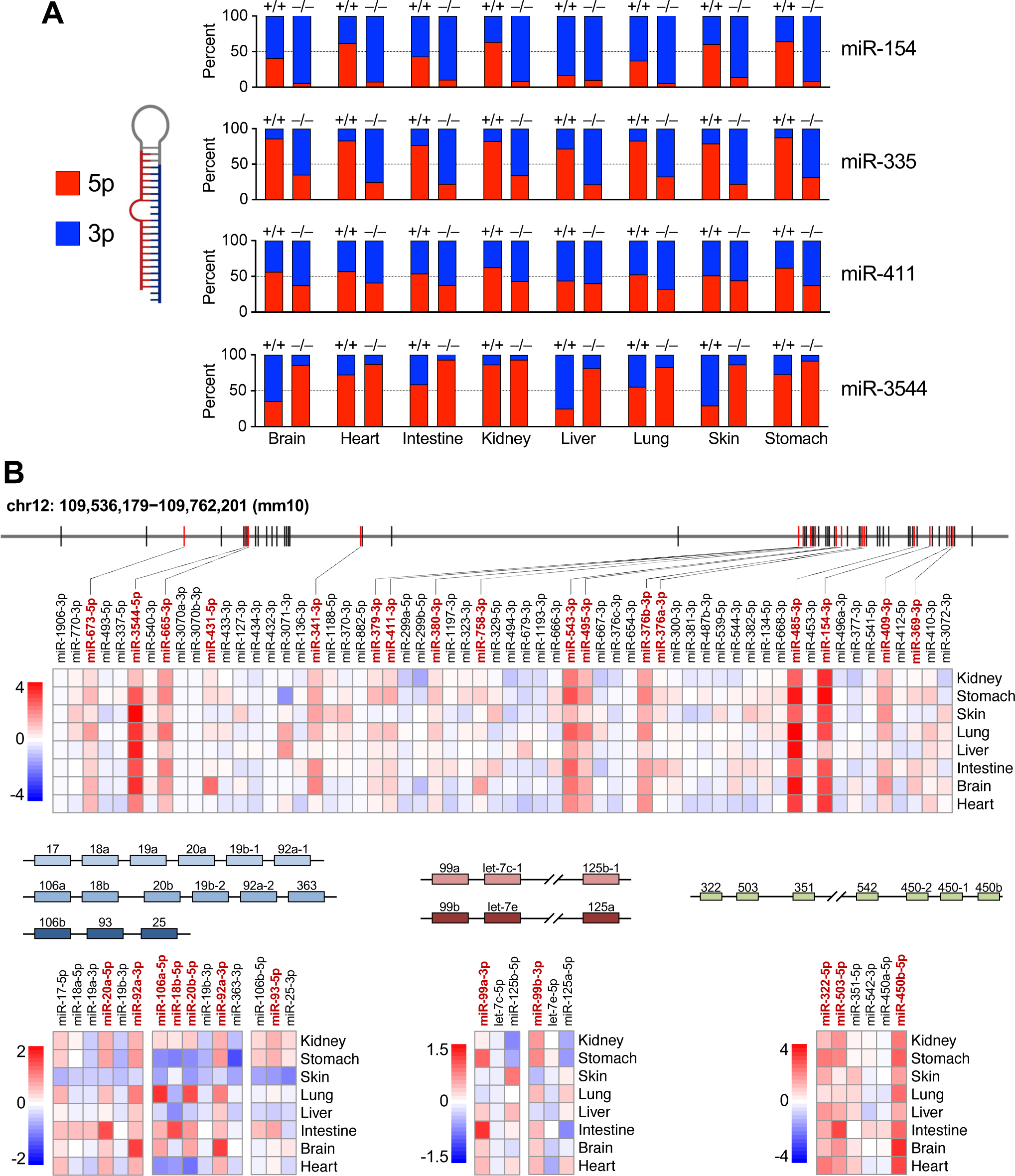
TDMD regulates non-dominant miRNA strands, arm switching, and clustered miRNAs. (*A*) Stacked bar graphs showing the relative abundance of 5p and 3p strands of the indicated miRNAs in E18.5 tissues from *Zswim8*^+/+^ or *Zswim8*^‒/‒^ mice. (*B*) Schematic representation of miRNA clusters encoding TDMD-regulated miRNAs. Heat maps display log_2_ fold change of miRNA expression (*Zswim8*^‒/‒^/*Zswim8*^+/+^) for each cluster member across tissues. miRNAs labeled in red text are TDMD substrates in at least one tissue.

Another interesting feature of mammalian TDMD substrates is their organization in clusters. While approximately 25% of all miRNAs are encoded in clusters (Kabekkodu et al. 2018), we observed that 36 out of 57 TDMD-regulated miRNAs (63%) were organized in this manner. This enrichment for clustered miRNAs is more than expected by chance (p<0.0001, chi-squared test). In total, we detected members of 14 different miRNA clusters that were regulated TDMD, notably including miRNAs encoded in a very large imprinted cluster on mouse chromosome 12 and members of the oncogenic miR-17-92 and paralogous clusters, among others (**Fig. 4B** and **Supplemental Fig. S3**). In some cases, individual cluster members were regulated by TDMD across all examined tissues, such as miRNAs derived from the imprinted chromosome 12 cluster and the miR-322-450b cluster (**Fig. 4B**). In other instances, clustered miRNAs were only regulated by TDMD in specific tissues, as exemplified by members of the miR-466 cluster (**Supplemental Fig. S3**). Considering that clustered miRNAs are generally co-transcribed, these results demonstrate that the TDMD pathway is employed as a means to post-transcriptionally regulate the expression of individual miRNAs within clusters in a tissue-specific manner.

It has been shown that extensive complementarity between the 3′ region of a miRNA and a TDMD-inducing target can stimulate the addition or removal of nucleotides from the miRNA 3′ end, a process known as tailing and trimming (Ameres et al. 2010). Although tailing and trimming is not required for TDMD (Han et al. 2020; Shi et al. 2020), its association with this pathway has been used as a feature to identify target transcripts that promote TDMD (Li et al. 2021). Moreover, in cell lines, loss of ZSWIM8 can increase the abundance of tailed and trimmed miRNAs that are normally regulated by TDMD (Han et al. 2020; Shi et al. 2020), presumably by stabilizing the interaction between miRNAs and TDMD-inducing targets that leads to modification of the miRNA 3′ end. Our northern blot analyses confirmed that a subset of TDMD-regulated miRNAs in mouse tissues undergo tailing and trimming (**Fig. 3C** and **Supplemental Fig. S2**). To examine this phenomenon more globally, the proportion of tailed or trimmed miRNA species in small RNA sequencing data from *Zswim8*^+/+^ and *Zswim8*^‒/‒^ mice was assessed. We observed a wide range of tailing and trimming of TDMD substrates in all tissues, with some miRNAs exhibiting negligible 3′ modification (**Supplemental Fig. S4A-B**). Loss of ZSWIM8 resulted in an increase in tailing and trimming of TDMD-regulated miRNAs compared to non-TDMD substrates in most tissues, with the exception of small intestine and lung where increased tailing was not observed (**Fig. 5A-B**). In half of the examined tissues, the magnitude of regulation of a miRNA by TDMD was positively correlated with its increased tailing and/or trimming upon loss of ZSWIM8 (**Fig. 5C** and **Supplemental Fig. S4C**). Overall, our data confirmed that tailing and trimming is a detectable feature of many TDMD substrates in vivo, but its prevalence varies across tissues, perhaps reflecting variation in the expression of the relevant terminal transferases or exoribonucleases. The fact that a subset of TDMD-regulated miRNAs do not undergo tailing and trimming may indicate that some miRNAs engage in alternative base-pairing configurations with their cognate TDMD-inducing triggers that do not expose the miRNA 3′ end.

**Figure 5.**
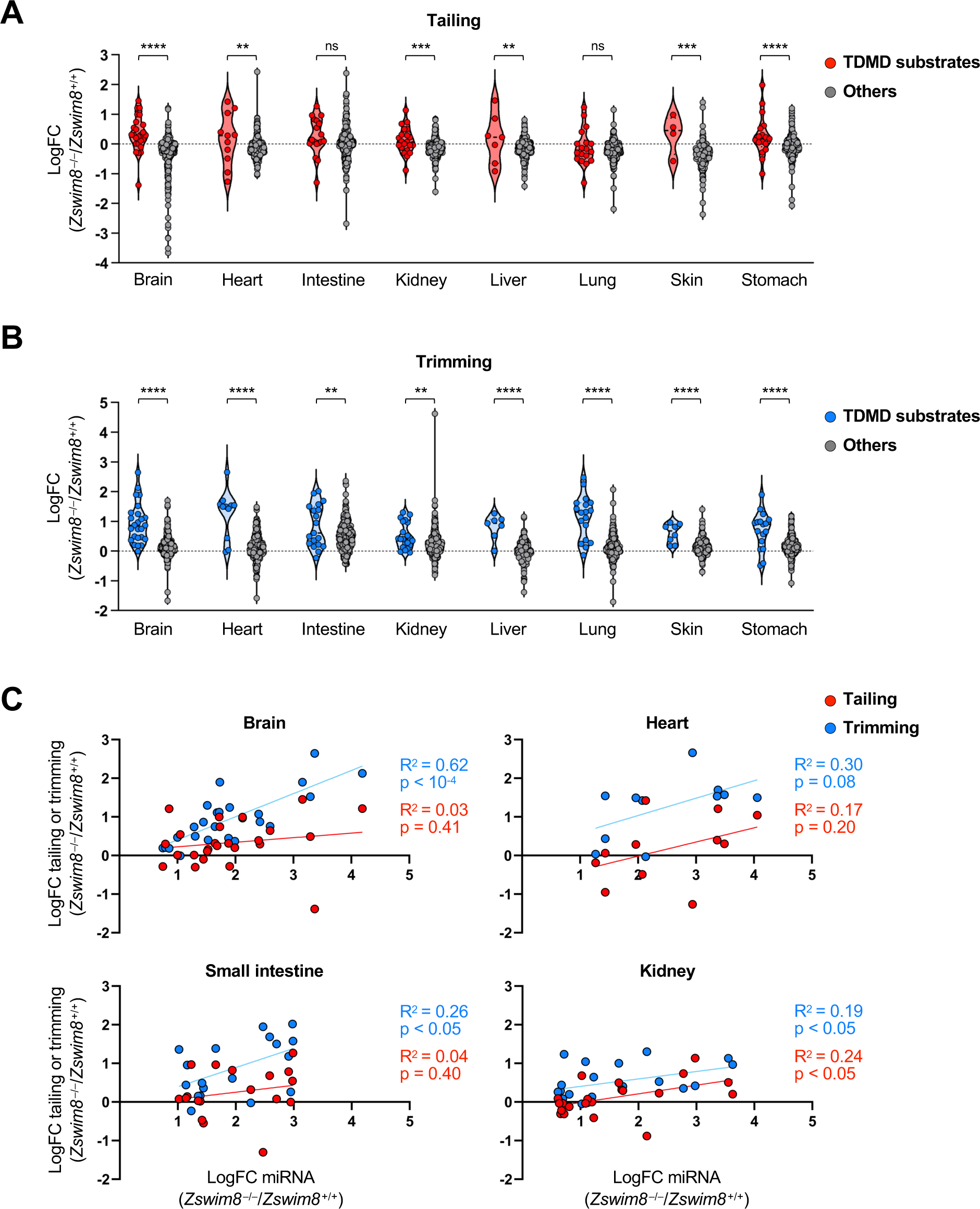
Tailing and trimming of TDMD-regulated miRNAs in mouse tissues. (*A*,*B*) Violin plots showing the fold change (*Zswim8*^‒/‒^/*Zswim8*^+/+^) of tailing (*A*) or trimming (*B*) of miRNAs in E18.5 tissues. ns, not significant; ****p<0.0001; ***p<0.001; **p<0.01 (unpaired *t*-test). (*C*) Scatter plots showing the fold change of tailing or trimming of each TDMD-regulated miRNA relative to its fold change in abundance (*Zswim8*^‒/‒^/*Zswim8*^+/+^) in each tissue.

### Growth restriction of *Zswim8^‒/‒^* mice is attributable to upregulation of miR-322 and miR-503

Among the most abundant miRNAs that were regulated by TDMD across multiple tissues were miR-322 and miR-503 (**Fig. 3A-B**), which are co-transcribed as part of a larger cluster of miRNAs located on the X chromosome (**Fig. 4B**). These miRNAs have previously been identified as TDMD substrates in murine cell lines (Rissland et al. 2011; Shi et al. 2020). As members of the miR-15/16 family, a highly-studied group of miRNAs that regulate the cell cycle and apoptosis (Aqeilan et al. 2010), we wondered whether dysregulation of miR-322 and miR-503 contributed to the developmental abnormalities observed in *Zswim8*^‒/‒^ mice. To investigate this possibility, Cas9 and dual sgRNAs were used to generate mice harboring a genomic deletion encompassing these miRNAs (**Fig. 6A**). As reported previously (Llobet-Navas et al. 2014), loss of these miRNAs had no effect on embryonic viability and mice carrying the knockout allele were present at Mendelian ratios at weaning (**Supplemental Fig. S5A-B**). Northern blotting confirmed the complete loss of expression of miR-322 and miR-503 in knockout tissues (**Fig. 6B**).

**Figure 6.**
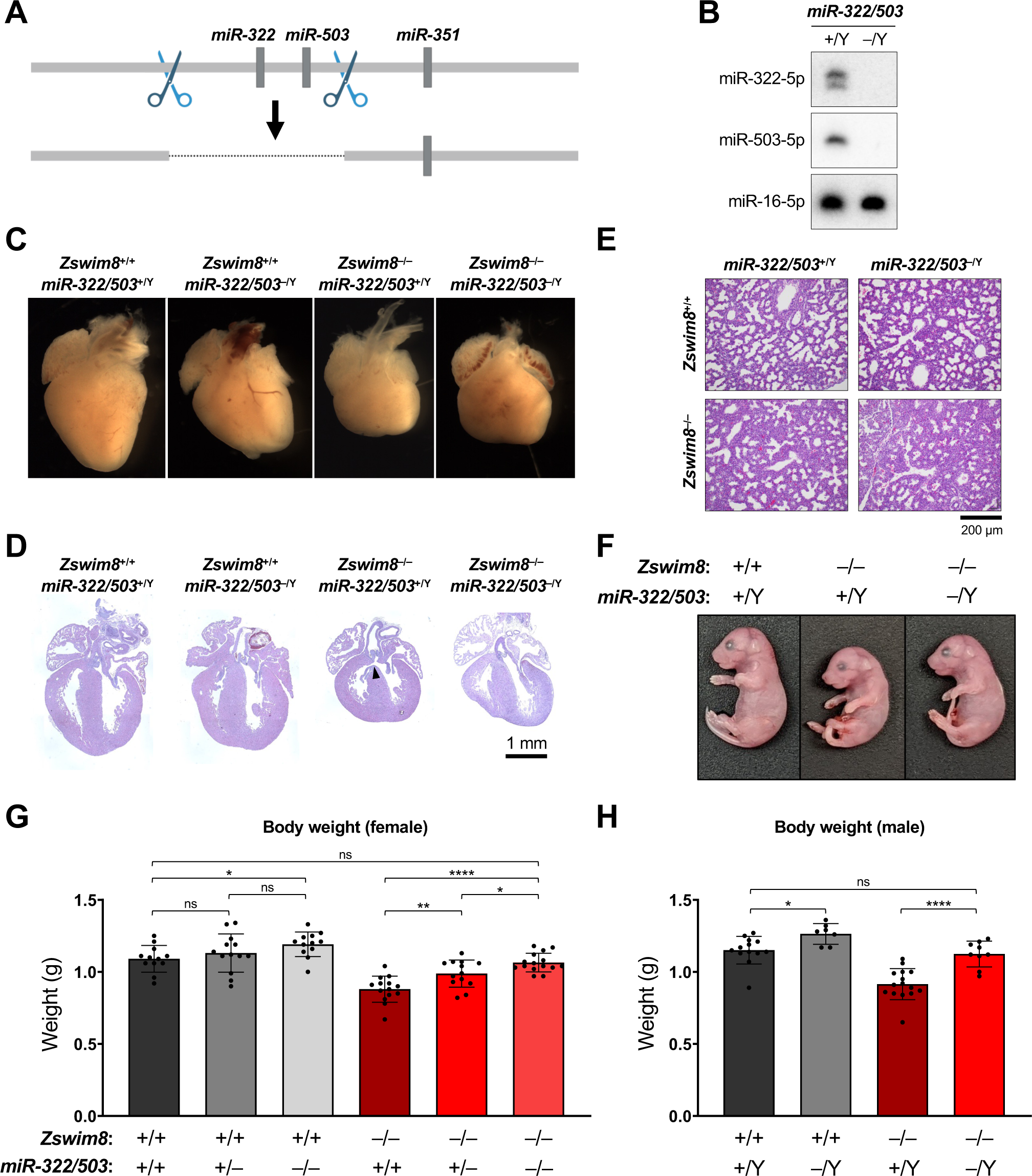
Loss of miR-322 and miR-503 rescues embryonic growth in *Zswim8^‒/‒^* mice. (*A*) Genome editing strategy used to generate *miR-322/503*^‒/‒^ mice. Scissors depict approximate location of CRISPR-Cas9 targeting sites. (*B*) Northern blot analysis of miRNA expression in E18.5 hearts of the indicated genotypes. (*C*) Images of E18.5 hearts of the indicated genotypes. (*D*,*E*) H&E-stained sections of E18.5 hearts (*D*) and lungs (*E*) of the indicated genotypes. VSD in *Zswim8*^‒/‒^; *miR-322/503*^+/Y^ heart in (D) indicated with arrowhead. (*F*) Images of E18.5 mice of the indicated genotypes. (*G*,*H*) Body weights of E18.5 female (*G*) and male (*H*) mice of the indicated genotypes. n = 7-15 mice per genotype. Data are represented as mean ± SD with individual data points shown. ns, not significant; ****p<0.0001; **p<0.01; *p<0.05 (unpaired *t*-test).

Mice carrying null alleles of *miR-322/503* and *Zswim8* were then intercrossed to generate double knockout animals. As observed in *Zswim8* single knockout mice, female *Zswim8*^‒/‒^; *miR-322/503*^‒/‒^ and male *Zswim8*^‒/‒^; *miR-322/503*^‒/Y^ mice died shortly after birth and were not present at weaning (**Supplemental Fig. S5A-B**). In line with the observed perinatal lethality of these animals, E18.5 double knockout mice exhibited malformed, globose hearts, with poorly defined apices and VSDs, as well as immature lungs at the late canalicular stage (**Fig. 6C-E** and **Supplemental Fig. S5C**). In contrast to these similarities between *Zswim8* single knockout mice and *Zswim8*; *miR-322/503* double knockouts, we observed that loss of miR-322 and miR-503 was sufficient to rescue embryonic growth of ZSWIM8-deficient mice (**Fig. 6F-H**). Analysis of body weights of E18.5 female mice enabled determination of the contribution of individual *miR-322/503* alleles to embryonic body size. Haploinsufficiency of *miR-322/503* resulted in significantly larger *Zswim8*^‒/‒^ embryos but did not significantly impact body weight in *Zswim8*^+/+^ mice (**Fig. 6G**). Homozygous loss of these miRNAs led to a full normalization of body size in the *Zswim8*^‒/‒^ background while also significantly increasing the growth of wild-type embryos. In males, where only complete absence of miR-322 and miR-503 could be examined since these miRNAs are encoded on the X chromosome, we again observed that deletion of *miR-322/503* increased body size in *Zswim8*^+/+^ animals but had a stronger effect in *Zswim8*^‒/‒^ mice, fully rescuing body size in this setting (**Fig. 6H**). Together, these results demonstrated that the *miR-322/503* locus is a dose-dependent negative regulator of embryonic growth and revealed that upregulation of the encoded miRNAs contributed to the small body size phenotype of *Zswim8*^‒/‒^ mice. These data therefore established the TDMD pathway as a regulator of embryonic growth in mammals.

## Discussion

TDMD was initially discovered as a mechanism of miRNA decay that can be induced by exogenously introduced targets, including transcripts encoded by viruses and synthetic targets with extensive miRNA complementarity (Ameres et al. 2010; Cazalla et al. 2010). The subsequent identification of a select group of endogenous miRNA:target pairs that trigger TDMD demonstrated that this pathway is a physiologic mechanism used to control miRNA abundance, although the number of known TDMD substrates remained limited (Bitetti et al. 2018; Ghini et al. 2018; Kleaveland et al. 2018; Li et al. 2021). The recent discovery of the ZSWIM8 ubiquitin ligase, which is essential for miRNA decay by TDMD (Han et al. 2020; Shi et al. 2020), has now enabled the direct experimental analysis of the scope and biological role of this pathway in diverse metazoan species (Kingston et al. 2022). Here we report the phenotypic consequences of ZSWIM8 loss of function in mice and characterize the landscape of TDMD-regulated miRNAs in mammalian tissues. These studies revealed that the TDMD pathway is required for normal mammalian development and is essential for establishing the appropriate expression of a broad set of miRNAs in vivo.

Global loss of ZSWIM8 resulted in an array of developmental defects in mice, including malformed hearts, immature lungs, and embryonic growth restriction, ultimately leading to fully-penetrant perinatal lethality. Nevertheless, the survival of *Zswim8*^‒/‒^ mice to birth enabled a comprehensive analysis of the repertoire of TDMD-regulated miRNAs in a broad set of late embryonic tissues. These studies revealed several notable features of the mammalian TDMD pathway. First, we detected 57 miRNAs that behaved as TDMD substrates in at least one tissue, as defined by a statistically-significant increase in the abundance of the miRNA in *Zswim8*^‒/‒^ mice without an increase in the opposite strand derived from the same precursor. Prior to this study, two mammalian miRNAs were identified as TDMD substrates in mouse tissues (Bitetti et al. 2018; Kleaveland et al. 2018) and another 32 miRNAs were shown to be regulated by this pathway in mammalian cell lines (Ghini et al. 2018; Han et al. 2020; Shi et al. 2020; Li et al. 2021). Our results therefore considerably expand the known catalog of TDMD-regulated miRNAs and suggest that this miRNA turnover pathway may be more active in vivo compared to cell lines. We speculate that this increased repertoire of TDMD substrates may reflect a need to efficiently clear specific miRNAs during rapidly-timed developmental transitions that are not modeled in cultured cells where mammalian TDMD has been previously studied. Additionally, it may be advantageous to regulate miRNAs via TDMD in terminally-differentiated, non-dividing cells in somatic tissues since, in contrast to cell lines, miRNAs cannot be diluted by cell division upon transcriptional shut-off in this setting. A second major feature of TDMD-regulated miRNAs in mammals is their frequent organization in genomic clusters. 36 out of the 57 TDMD-regulated miRNAs are derived from clusters, representing significant enrichment. Since clustered miRNAs are generally co-transcribed, TDMD appears to be regularly employed as a mechanism to uncouple the expression of individual cluster members. Third, we identified examples of miRNAs in which the non-dominant strand was strongly regulated by TDMD and, in some cases, became the dominant strand in specific ZSWIM8-deficient tissues. This observation suggests that some examples of the previously described phenomenon of ‘arm switching’, in which the dominant miRNA strand from a miRNA precursor varies across cell types or conditions, may be due to selective decay of one miRNA strand by the TDMD pathway. Indeed, we found that tissue-specific differences in the magnitude of downregulation of the 3p strand of miR-154 by TDMD provides a mechanistic explanation for the reported arm switching of this miRNA across tissues (Cheng et al. 2013). Altogether, these data illuminate the broad scope of TDMD in mammalian tissues and reveal previously unrecognized roles for this pathway in shaping miRNA expression in vivo.

The phenotypic consequences of ZSWIM8 loss of function vary across metazoan species. In *C. elegans,* loss of the ZSWIM8 ortholog EBAX-1 results in neuromuscular defects, reduced egg laying, and decreased male mating (Wang et al. 2013). Deficiency of Dora, the ZSWIM8 ortholog in *Drosophila*, leads to embryonic lethality and defective cuticle development (Kingston et al. 2022). As reported here, mouse ZSWIM8 is essential for normal heart and lung development, embryonic growth, and post-natal viability. However, as a component of a ubiquitin ligase complex, ZSWIM8 may regulate the stability of multiple proteins in addition to AGO, thus complicating the interpretation of these phenotypes. Indeed, ZSWIM8 homologs in *C. elegans* and mouse have been reported to impact neurodevelopmental phenotypes by controlling the stability of proteins involved in axon guidance and brain development (Wang et al. 2013; Wang et al. 2023). Thus far, the strongest evidence that the TDMD pathway itself plays an important role in regulating miRNA expression during development comes from studies of *Drosophila*. Reduction of the dose of the TDMD-regulated miRNAs miR-3 and miR-309 partially rescued embryonic lethality in Dora-mutant flies and loss of the target transcript that triggers TDMD of the miR-310 family impaired cuticle development (Kingston et al. 2022). Thus, in *Drosophila*, regulation of miRNA expression by the TDMD pathway is essential for normal development. Our findings reported here now extend this principle to mammals. Specifically, we demonstrated that reduced dosage of the TDMD-regulated miRNAs miR-322 and miR-503 rescued embryonic growth in ZSWIM8-deficient mice. Loss of a single allele of the *miR-322/503* locus significantly increased body size in *Zswim8*^‒/‒^ embryos without affecting growth of wild-type embryos, while complete loss of these miRNAs increased body size in all genotypes. These results demonstrate that miR-322 and miR-503 function as dose-dependent regulators of embryonic growth and the strongly elevated expression of these miRNAs in ZSWIM8-deficient embryos plays a causative role in the growth restriction phenotype. While the specific targets of miR-322 and miR-503 that mediate their growth-suppressive effects remain to be determined, the previously established roles for these miRNAs in regulation of the cell cycle and insulin-like growth factor signaling are promising directions for future investigation given the central roles of these pathways in mammalian body size regulation (Conlon and Raff 1999; Linsley et al. 2007; Rissland et al. 2011; Llobet-Navas et al. 2014). Further studies in which the dosage of other TDMD-regulated miRNAs is reduced in ZSWIM8-deficient mice, or TDMD-inducing target sites are ablated, will be important to determine whether miRNA dysregulation underlies defective heart and lung development in *Zswim8*^‒/‒^ embryos.

Looking ahead, a major challenge for the future will be identifying the target sites that trigger decay of the broad set of TDMD-regulated miRNAs described here. Only four endogenous mammalian TDMD triggers have been identified to date (Bitetti et al. 2018; Ghini et al. 2018; Kleaveland et al. 2018; Li et al. 2021), leaving dozens left to be discovered. In order to address this deficit, all modes of target:miRNA interaction that are capable of triggering TDMD must be determined. All known examples of TDMD triggers in mammals exhibit extensive complementarity to both the seed region and 3′ end of the miRNA. Recent work from *C. elegans*, however, has demonstrated that the ZSWIM8 ortholog EBAX-1 can promote the degradation of specific miRNAs in a manner depending only on the seed sequence (Donnelly et al. 2022). This raises the possibility of alternative base-pairing configurations between some miRNAs and their TDMD-inducing targets in mammals. Interestingly, tailing and trimming of miRNAs has been associated specifically with extensive 3′ complementarity to the target, which removes the miRNA 3′ end from the AGO PAZ domain and exposes it to the activity of terminal transferase and exonuclease enzymes (Ameres et al. 2010; Sheu-Gruttadauria et al. 2019; Yang et al. 2020). While many mammalian TDMD substrates exhibit tailing and trimming, some do not, raising the possibility that this latter class of miRNAs may be targeted for TDMD by alternative base-pairing configurations. In these cases, biochemical methods such as crosslinking and immunoprecipitation (CLIP) (Lee and Ule 2018), which could identify RNAs associated with ZSWIM8, might be informative for target identification. Further identification of TDMD triggers and their substrates will be critical for delineating the functions of this important miRNA regulatory pathway in physiology and disease.

## Materials and Methods

### Generation and analysis of genetically engineered mice

All mouse experiments were approved by University of Texas Southwestern Medical Center Animal Care and Use Committee and performed in accordance with NIH guidelines (Animal protocol 2017-102001). Mice were group housed under conventional conditions in a 12-hour day/night cycle with ad libitum availability of normal chow diet (Harlan Teklad, TD2916) and water. *Nkx2.5*-Cre mice were generously provided by Drs. Eric Olson and Rhonda Bassel-Duby (UT Southwestern).

*Zswim8*^‒/‒^, *miR322/503*^‒/‒^, and *Zswim8*^F/F^ mice were generated using CRISPR/Cas9-mediated genome editing at the UT Southwestern Transgenic core. To generate *Zswim8*^‒/‒^ and *miR322/503*^‒/‒^ mice, Cas9 protein in complex with sgRNAs (Integrated DNA Technologies) were either microinjected or electroporated into the cytoplasm of fertilized C57BL/6J fertilized eggs. To generate *Zswim8*^F/F^ mice, loxP-containing Megamer oligos (Integrated DNA Technologies) were co-injected with Cas9-sgRNA complexes into the pronucleus of fertilized C57BL/6 eggs. Sequences of sgRNAs and megamer are provided in **Supplemental Table S2**. Founders carrying the desired alleles were maintained in a pure C57BL/6J background with continuous backcrossing. For timed matings, the morning of detection of vaginal plug was defined as E0.5.

Phenotyping of embryos was performed at E18.5 after delivery by caesarean section. Embryos were weighed and imaged before sacrificing. Tissues were fixed in 10% neutral-buffered formalin (Sigma) overnight, followed by paraffin embedding and sectioning for H&E staining. Tissue sections were imaged using a Zeiss Axio Observer Z1 microscope and whole mount images were generated using a Nikon SMZ800N with NIS Elements software (Nikon). High resolution composite images of the entire section (heart and lung) were captured using tiling and stitching functions of Zen Pro 3.4 software (Blue edition; Zeiss).

### Western blotting

Brains from E18.5 mice were homogenized in lysis buffer (40 mM HEPES, pH 7.4, 2 mM EDTA, 2 mM EGTA, 150 mM NaCl, 1% Triton X-100) supplemented with 1X complete protease inhibitor cocktail and 1X complete phosphatase inhibitor cocktail (Roche) using a Precellys Evolution homogenizer (Bertin Technologies). Approximately 100 μg of lysate was resolved on 8% SDS-PAGE, blotted onto nitrocellulose membranes, and probed with anti-ZSWIM8 (1:400; Cat #PA5-59492, Thermo) or anti-GAPDH (1:1000; Cat #2118, Cell Signaling) followed by HRP-conjugated anti-rabbit secondary (1:10,000; Cat #111-035-144, Jackson ImmunoResearch). Chemiluminescence detection was performed using Femto Clean ECL Solution (Thermo Fisher) with a Chemidoc MP Imaging Station (BioRad).

### Quantitative PCR measurement of floxed allele recombination

E18.5 hearts were homogenized using a Precellys Evolution homogenizer (Bertin Technologies) and genomic DNA was extracted using the DNeasy Blood and Tissue kit (Qiagen). Quantitative PCR was performed with Power SYBR Green PCR Master Mix (Thermo Fisher) using primers that were specific for the non-recombined allele. Primers that amplified a region of the *Zswim8* gene outside of the targeted sequence were used for normalization. Percent recombined was calculated as 1−(frequency of non-recombined alleles). Primer sequences provided in **Supplemental Table S2.**

### RNA extraction, small RNA sequencing, and northern blot analysis

Tissues from E18.5 embryos were homogenized in Qiazol (Qiagen) using a Precellys Evolution homogenizer (Bertin Technologies). Total RNA was extracted using the miRNeasy Mini RNA isolation kit (Qiagen) and digested with DNase I to remove genomic DNA. Library preparation was performed as previously described (Kim et al. 2019; Han et al. 2020). In brief, total RNA was size fractionated on a 15% urea-polyacrylamide gel. RNAs were ligated to a 3¢ randomized adapter using T4 RNA ligase 2 truncated KQ in buffer supplemented with 20% PEG 8000 at 25°C overnight. Following gel-purification, RNAs were ligated to a 5¢ randomized adapter using T4 ligase 1 in buffer supplemented with 20% PEG 8000 at 37°C for one hour. The product was then reverse transcribed with Superscript III (Invitrogen), cDNA was PCR amplified with Phusion polymerase (Thermo Fisher), and product was gel purified.

Northern blotting was performed as previously described (Han et al. 2020). In brief, total RNA was run on a 15% urea-polyacrylamide gel and transferred to a nylon membrane before UV-crosslinking, blocking, and probing with a radiolabeled probe. A locked nucleic acid probe (mmu-miR-450b-5p miRCURY LNA miRNA Detection probe; Qiagen) was used to detect miR-450b while standard DNA oligonucleotide probes were used for other miRNAs. Probe sequences provided in **Supplemental Table S2**.

### Analysis of small RNA sequencing data

Adapters were trimmed using cutadapt (Martin 2011), and trimmed reads with low quality (-Q33 -q 20 -p 95) were filtered out using FASTX-Toolkit (http://hannonlab.cshl.edu/fastx_toolkit/). Reads were assigned to miRNAs in miRbase v22 (Kozomara et al. 2019) based on a perfect match to the first 18 nucleotides of each miRNA. Tailing and trimming was analyzed as previously described (Kleaveland et al. 2018). For each miRNA, the proportion of reads with each observed length was calculated. The minimum length of the normal range (*L_min_*) of a miRNA was set as the minimum length with a proportion greater than 14.5%, and the maximum length of the normal range (*L_max_*) was set as the maximum length with a proportion greater than 14.5%. The reads with length between *L_min_* and *L_max_* were defined as non-tailed/non-trimmed miRNA, the reads with length less than *L_min_* were defined as trimmed miRNA, and reads with length greater than *L_max_* were defined as tailed miRNA. EdgeR (Robinson et al. 2010) was used to identify differentially expressed miRNAs between genotypes.

### Quantification and statistical analysis

For measurements of body weight, each data point represents a single embryo or mouse. Statistical comparisons were performed using Prism 9. Unpaired t-tests were performed to evaluate the difference between different genotypes, as well as the differences in tailing and trimming between miRNAs. A chi-square test was used to evaluate the enrichment of clustered miRNAs among TDMD substrates. Significance is indicated as follows: *p<0.05, **p<0.01, ***p<0.001, ****p<0.0001. All values are reported as mean ± SD.

## Acknowledgements

We thank Eric Olson and Rhonda Bassel-Duby for mouse strains; Mylinh Nguyen in the UT Southwestern Transgenic Core; John Shelton in the UT Southwestern Histo Pathology Core; Vanessa Schmid in the McDermott Center Next Generation Sequencing Core; Jeanetta Marshburn-Wynn for assistance with mouse husbandry; and Kathryn O’Donnell and members of the Mendell laboratory for helpful suggestions on the manuscript. This work was supported by grants from CPRIT (RP220309 to J.T.M.), the Welch Foundation (I-1961-20210327 to J.T.M.), and NIH (R01CA282036 to J.T.M.). J.T.M. is an Investigator of the Howard Hughes Medical Institute. This article is subject to HHMI’s Open Access to Publications policy. HHMI lab heads have previously granted a nonexclusive CC BY 4.0 license to the public and a sublicensable license to HHMI in their research articles. Pursuant to those licenses, the author-accepted manuscript of this article can be made freely available under a CC BY 4.0 license immediately upon publication.

## Author contributions

B.T.J., J.H., R.E.H., B.M.E., D.R., A.A., and J.T.M. designed experiments and interpreted results. B.T.J., J.H., and A.A. performed experiments. H.Z. and J.H. performed bioinformatic analyses. B.T.J., J.H., A.A., and J.T.M. wrote the manuscript.

