## Supplemental Figures for "Target-directed microRNA degradation regulates developmental microRNA expression and embryonic growth in mammals"

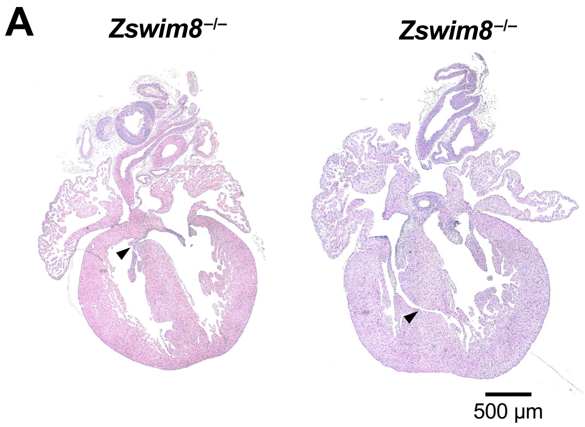


**B**

**P21**

| Genotype | Expected | Actual |
|---|---|---|
| *Zswim8*$^{+/+}$ | 15.125 | 20 |
| *Zswim8*$^{F/+}$ | 30.25 | 39 |
| *Zswim8*$^{F/F}$ | 15.125 | 18 |
| *Nkx2.5*-Cre; *Zswim8*$^{+/+}$ | 15.125 | 7 |
| *Nkx2.5*-Cre; *Zswim8*$^{F/+}$ | 30.25 | 28 |
| *Nkx2.5*-Cre; *Zswim8*$^{F/F}$ | 15.125 | 9 |

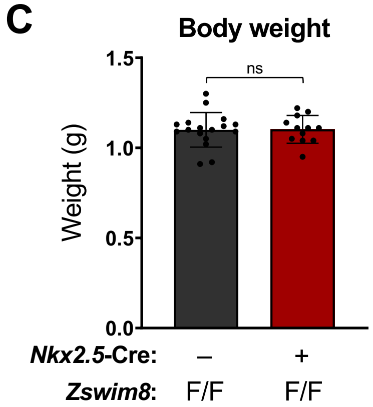


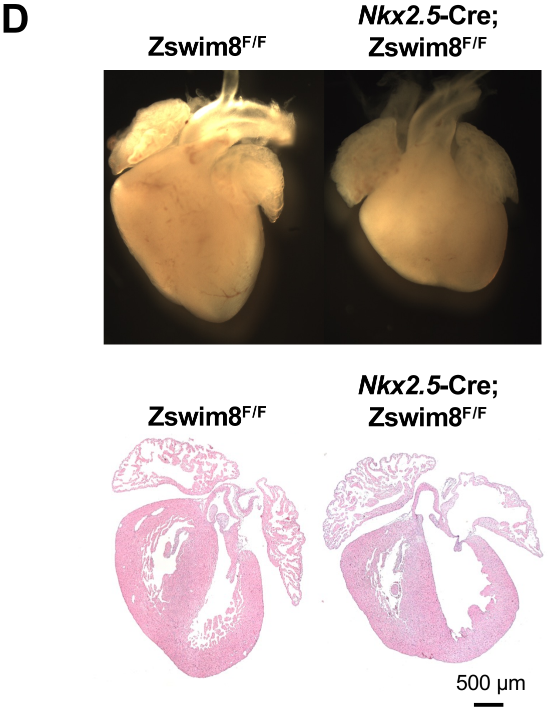


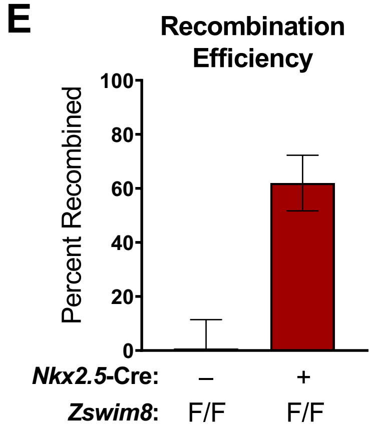


**Supplemental Figure S1. Analysis of mice with global or cardiac-specific loss of *Zswim8*.** (*A*) Hearts from E18.5 *Zswim8*$^{–/–}$ mice showing presence of VSDs (arrowheads). (*B*) Frequency of genotypes of offspring at P21 produced from crossing *Nkx2.5*-Cre; *Zswim8*$^{+/F}$ to *Zswim8*$^{+/F}$ mice. (*C*) Body weights of E18.5 mice of the indicated genotypes. n=17 (*Zswim8*$^{F/F}$) and 12 (*Nkx2-5*$^{Cre}$; *Zswim8*$^{F/F}$). Data are represented as mean ± SD with individual data points shown. ns, not significant (unpaired *t*-test). (*D*) Images and H&E-stained sections of E18.5 hearts of the indicated genotypes. (*E*) Quantitative PCR measurement of *Zswim8*$^{F/F}$ allele recombination efficiency in E18.5 hearts from mice of the indicated genotypes. Data are represented as mean ± SD.

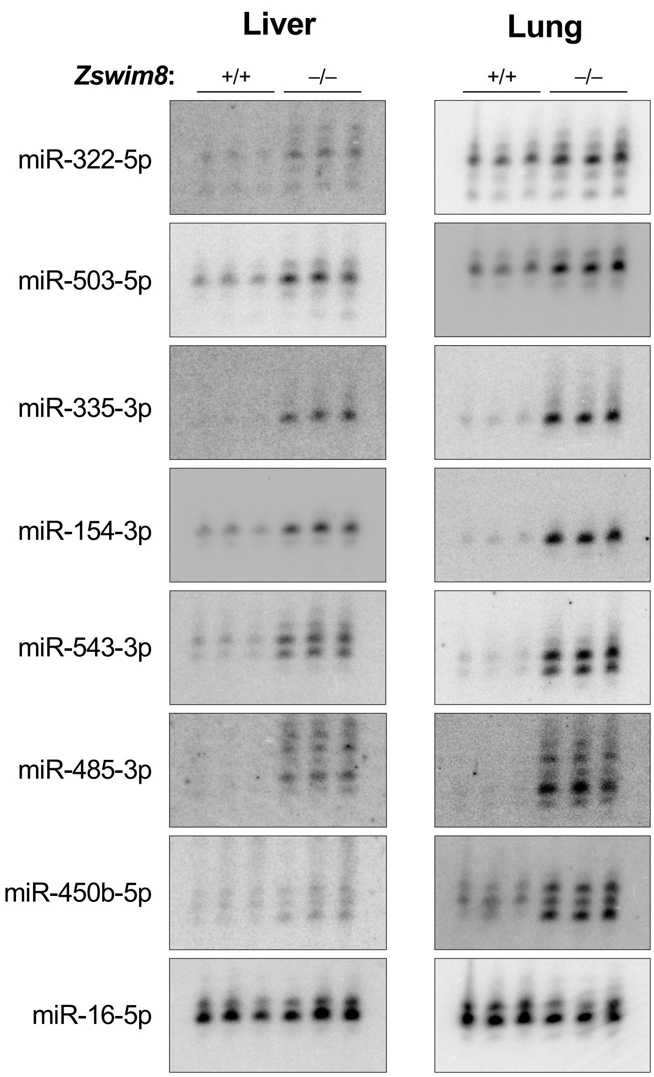


**Supplemental Figure S2**. **Northern blots of TDMD-regulated miRNAs in *Zswim8*$^{-/-}$ tissues.** Northern blot analysis of miRNA expression in E18.5 liver and lung from mice of the indicated genotypes. n=3 biological replicates per genotype. miR-16-5p served as a loading control.

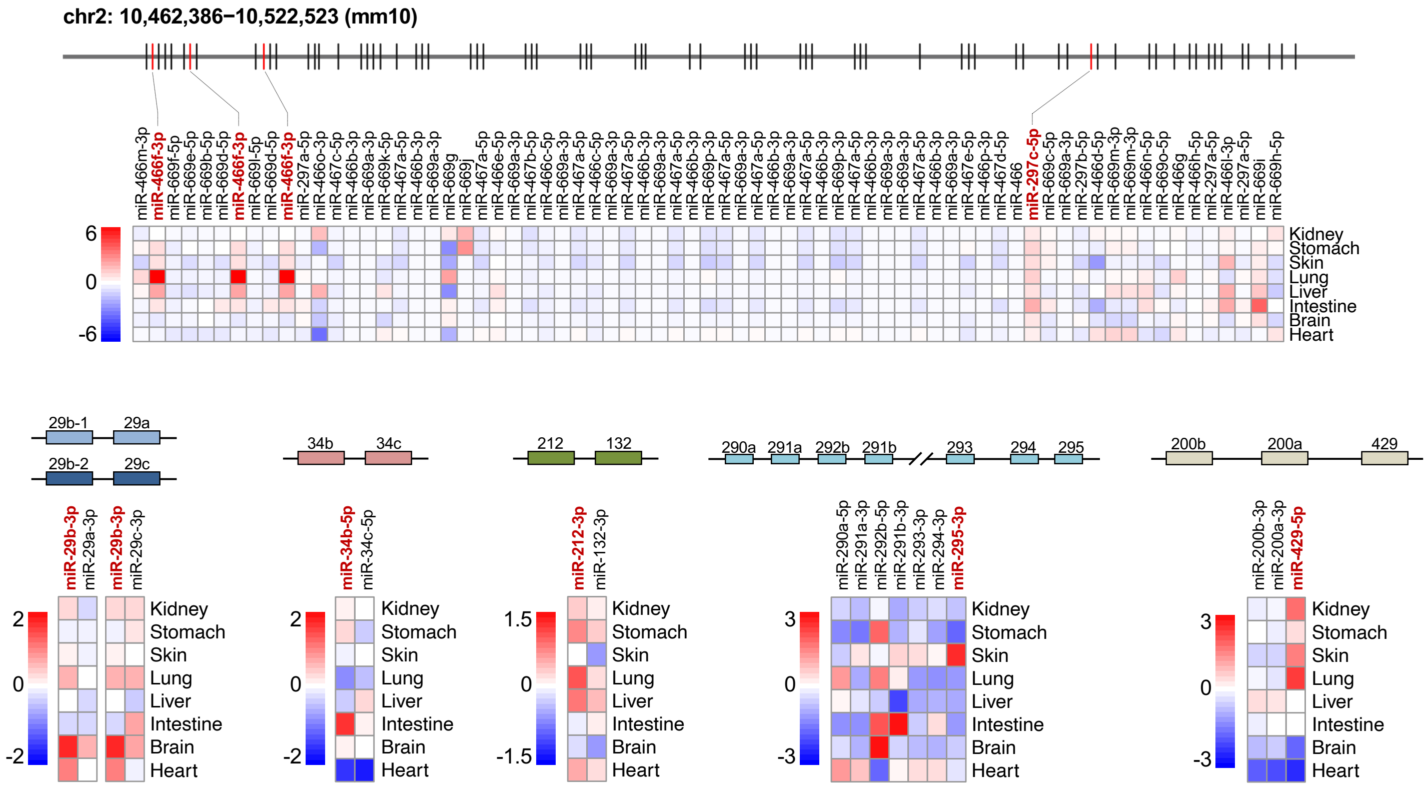


**Supplemental Figure S3**. **TDMD regulates clustered miRNAs.** Schematic representation of miRNA clusters encoding TDMD-regulated miRNAs. Heat maps display $\log_2$ fold change of miRNA expression (*Zswim8*$^{-/-}$/*Zswim8*$^{+/+}$) for each cluster member across tissues. miRNAs labeled in red text are TDMD substrates in at least one tissue.

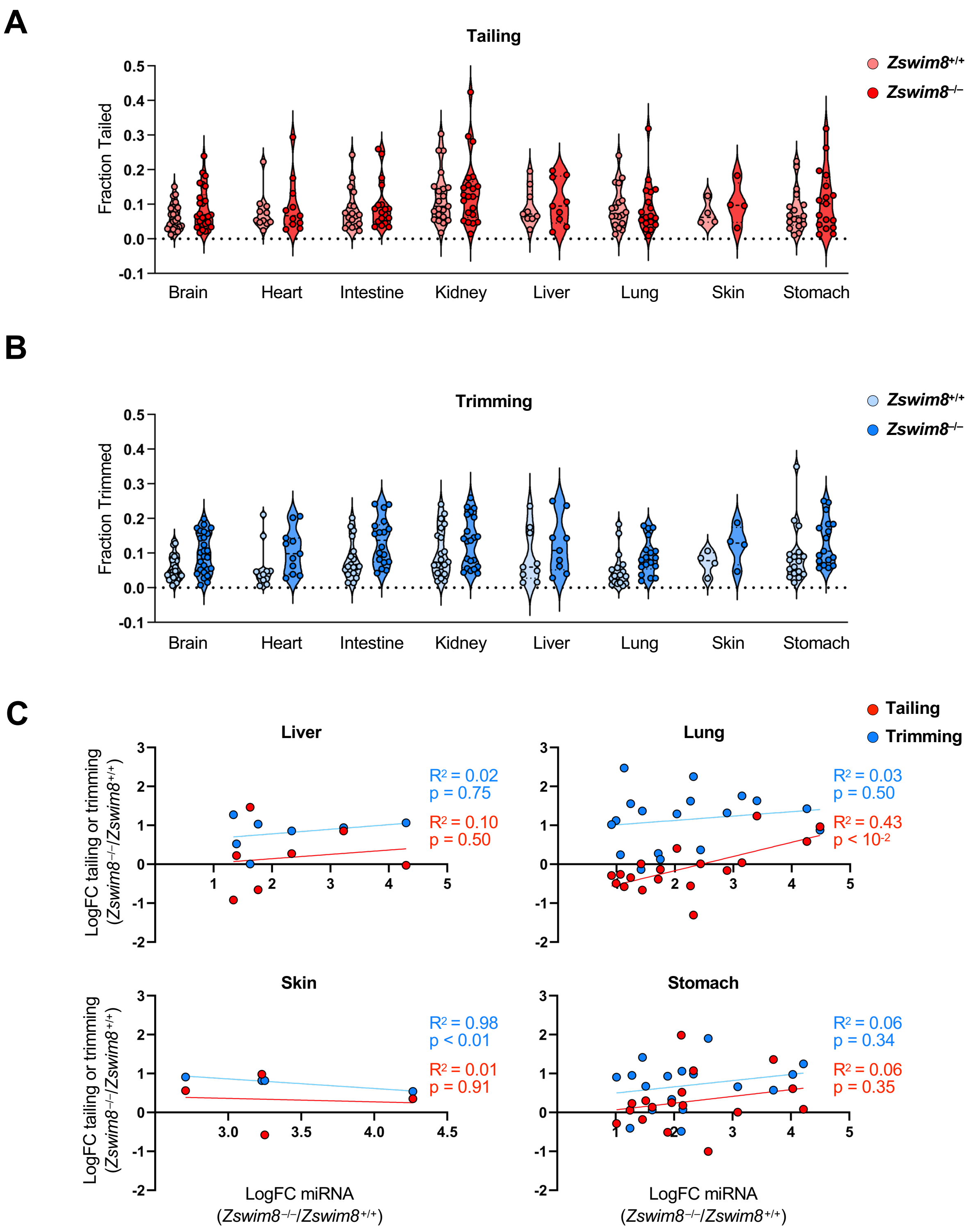


**Supplemental Figure S4. Tailing and trimming of TDMD-regulated miRNAs.** (*A*,*B*) Violin plots showing the fraction of tailed (*A*) or trimmed (*B*) isoforms of each TDMD-regulated miRNA in E18.5 tissues from *Zswim8*$^{+/+}$ or *Zswim8*$^{-/-}$ mice. (*C*) Scatter plots showing the fold change of tailing or trimming of each TDMD-regulated miRNA relative to its fold change in abundance (*Zswim8*$^{-/-}$/*Zswim8*$^{+/+}$) in each tissue.

**A**

**P21 (male)**

| ***Zswim8*** | ***miR-322/503*** | **Expected** | **Actual** |
|---|---|---|---|
| +/+ | +/Y | 7.125 | 13 |
| +/+ | –/Y | 7.125 | 8 |
| +/– | +/Y | 14.25 | 21 |
| +/– | –/Y | 14.25 | 15 |
| –/– | +/Y | 7.125 | 0 |
| –/– | –/Y | 7.125 | 0 |

**B**

**P21 (female)**

| ***Zswim8*** | ***miR-322/503*** | **Expected** | **Actual** |
|---|---|---|---|
| +/+ | +/– | 8.375 | 16 |
| +/+ | –/– | 8.375 | 13 |
| +/– | +/– | 16.75 | 20 |
| +/– | –/– | 16.75 | 18 |
| –/– | +/– | 8.375 | 0 |
| –/– | –/– | 8.375 | 0 |

**C**

***Zswim8*** $^{-/-}$
***miR-322/503*** $^{-/-}$

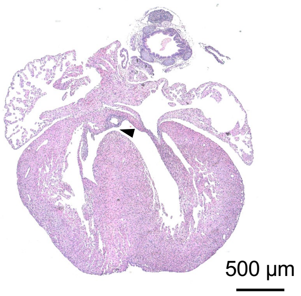


**Supplemental Figure S5**. **Loss of miR-322 and miR-503 does not rescue perinatal lethality or heart defects in *Zswim8*$^{-/-}$ mice.** (*A*,*B)* Frequency of genotypes of male (*A*) and female (*B*) offspring at P21 produced from intercrossing *Zswim8*$^{+/-}$; *miR-322/503*$^{+/-}$ females to *Zswim8*$^{+/-}$; *miR-322/503*$^{-/Y}$ males. (*C*) H&E-stained section of E18.5 *Zswim8*$^{-/-}$; *miR-322/503*$^{-/-}$ heart mice showing presence of VSD (arrowhead).
